# A serial dilution method for assessment of microplastic toxicity in suspension

**DOI:** 10.1101/401331

**Authors:** Zandra Gerdes, Markus Hermann, Martin Ogonowski, Elena Gorokhova

## Abstract

The occurrence of microplastic (MP) in the environment is of global concern. MP risk assessment, however, is currently hampered by lacking ecotoxicological methods due to conceptual and practical problems with particle exposure. Natural particles of similar size as MP, e.g., clay and cellulose, occur abundantly in the environment. For MP risk assessment and regulation it must be established whether the addition of MP to these particles represents an additional hazard. We present a novel approach employing a serial dilution of MP and reference particles, in mixtures, which allows the differentiation of MP effects from other particulates. We demonstrate the applicability of the method using an immobilisation test with *Daphnia magna* exposed to polyethylene terephthalate (MP) and kaolin clay (reference material). In the concentration range of 0.1 to 10000 mg L^-1^ of total suspended solids (TSS), with MP contributing 0-100 %, the LC_50_ values for MP-kaolin mixtures were significantly lower compared to the pure kaolin suspension. MP particles were thus more harmful to daphnids than the reference material. The estimated threshold for %MP contribution above which higher mortality was observed was 1 % MP at 36 mg TSS L^-1^. This approach has a potential for standardisation of MP ecotoxicological testing as well as other particulate material of anthropogenic origin.

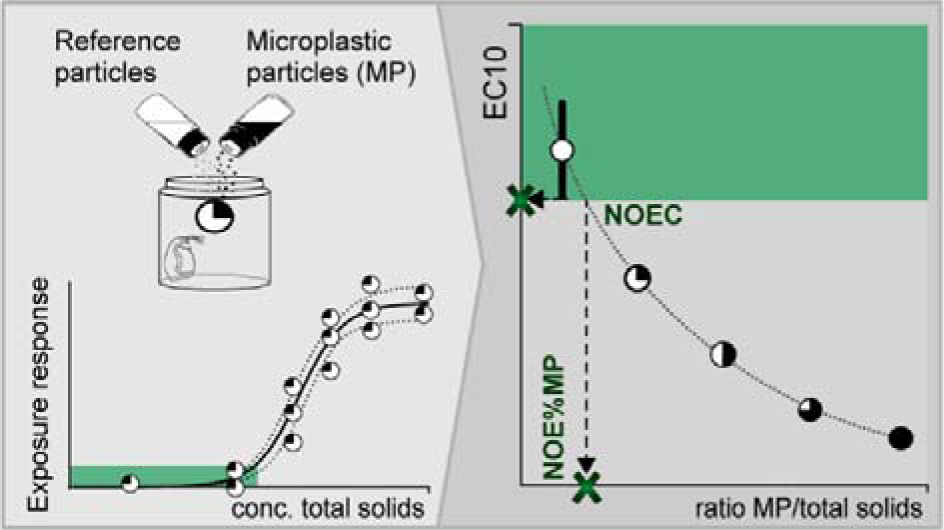

## Introduction

The increasing environmental pollution with plastic waste is of global concern. What is more, this debris eventually breaks down to small fragments collectively termed microplastics (MP) that are omnipresent in aquatic environments, including alpine lakes, rivers, oceans and arctic ice.^1–4^ The amounts of the plastic debris in general, and MP, in particular, are expected to increase because of increased production, continuous discharge, and fragmentation.^5^ Research on the hazard assessment of solid polymer particulates is in high demand due to public and scientific concerns. Nevertheless, scientists disagree on the immediacy of the MP pollution problem,^6–9^ and it remains largely unclear whether MP are harmful to biota and what the impact mechanisms are. The continuing uncertainty is, at least partly, related to the fact that MP are a new type of environmental contaminant with yet unsettled methodology for hazard testing.

The first experimental MP effect studies included a wide range of animal species focusing mainly on feeding-related impacts in filter-feeders, such as bivalves^10–12^ and zooplankton^13,14^ Filter-feeders continue to be among the commonly used test organisms in MP effect studies because they are susceptible to MP exposure via ingestion. Since MP particles are nutritionally inert, their ingestion decreases the energy intake. In other words, the ingestion of refractory material and alterations in feeding (a primary response) leads to lower growth and reproduction (secondary responses) as a result of the decreased caloric intake.^15^

All these processes occur not only with MP but also with any other refractory material present in natural seston. Both mineral^15–18^ and MP^13,14,19,20^ particles have been reported to alter feeding activity and reduce growth. Natural processes, such as wind and resuspension, primarily affect the presence of nutritionally inert particles in the water; whereas, human activities, like, dredging and stormwater runoff, may also elevate their concentrations. High concentrations of total suspended solids (TSS) have been found to reduce primary production,^21^ suppress population growth of zooplankton^22^ and alter feeding behaviour in fish.^23^ Therefore, to protect wildlife, water quality standards are implemented for TSS concentrations or allowable TSS levels in, e.g. stormwater effluents,^24^ lakes and streams^25^.

Regulatory efforts to set allowable MP levels are calling for adequate methodological approaches for hazard assessment, relevant model species, and exposure scenarios. A step towards quantifying hazardous properties of synthetic polymer microparticles is to develop and apply standardised practices and experimental designs that will be able to provide threshold values of these effects. However, given the presence of various particulates and the hazardous effects of high TSS concentrations, such designs should include the MP in question together with environmentally relevant reference material(s). Particular attention should be paid to the similarity of basic physical properties that are important for biological responses, e.g., size distribution and shape, between the reference particles and the MP.^13,26^ Also, to maintain the experimental reproducibility and stable encounter rates in a pelagic exposure scenario, it is important that all particles be kept in suspension during the incubation.

A recent comparison of the effects exerted by MP and mineral particulates suggests some similarity in responses across different levels of biological organisation, albeit with an indication of a greater hazard by MP.^27^ Since natural particles are more abundant than MP in aquatic environments,^7^ the hazardous levels of MP should rather be presented as a relative contribution of MP to TSS and not the absolute concentrations.

To date, there is no standard approach for MP effect assessment,^28^ despite a rapidly rising number of reports on MP effects under laboratory conditions. This is partly because it is challenging to design exposure experiments with environmentally relevant concentrations of MP based on the commonly reported levels (<10 particles m^3^)^29–31^. Moreover, the existing approaches do not explicitly test the effects of MP *per se* but those of nutrition-free particulates. To move the field of MP ecotoxicology forward, we need to use test methods that (1) are appropriate for delineating effects of different particulate materials in mixtures, (2) provide estimation of the critical concentrations of MP in different environments, (3) allow high-throughput testing, and (4) support read-across and categorical assessment of solid polymer particles. Here, we propose a new approach employing a linear serial dilution of MP and reference particles in mixtures, to identify MP-specific toxicity while controlling for the total concentration of suspended matter in the experimental system. Further, we demonstrate the applicability of this approach using the 96-h exposure of the cladoceran *Daphnia magna* to a mixture of polyethylene terephthalate (PET) as a test MP and kaolin as a reference particle.

## METHOD

The MP Ratio Test was designed to examine whether a particulate material (test particle) is harmful when co-occurring in a mixture with naturally present particulates (reference particle) across a range of TSS concentrations. The rationale is as follows: if the test particle is more harmful than the reference particle, then decreasing its contribution to a mixture with the reference particles should decrease the overall toxicity, assuming additivity of the effects (fig. 1). When the test (MP) and reference (mineral) particles are provided at varying proportions for each TSS concentration, then, by using a range of TSS concentrations, a dose-response relationship can be established for each mixture.

**Figure 1.**
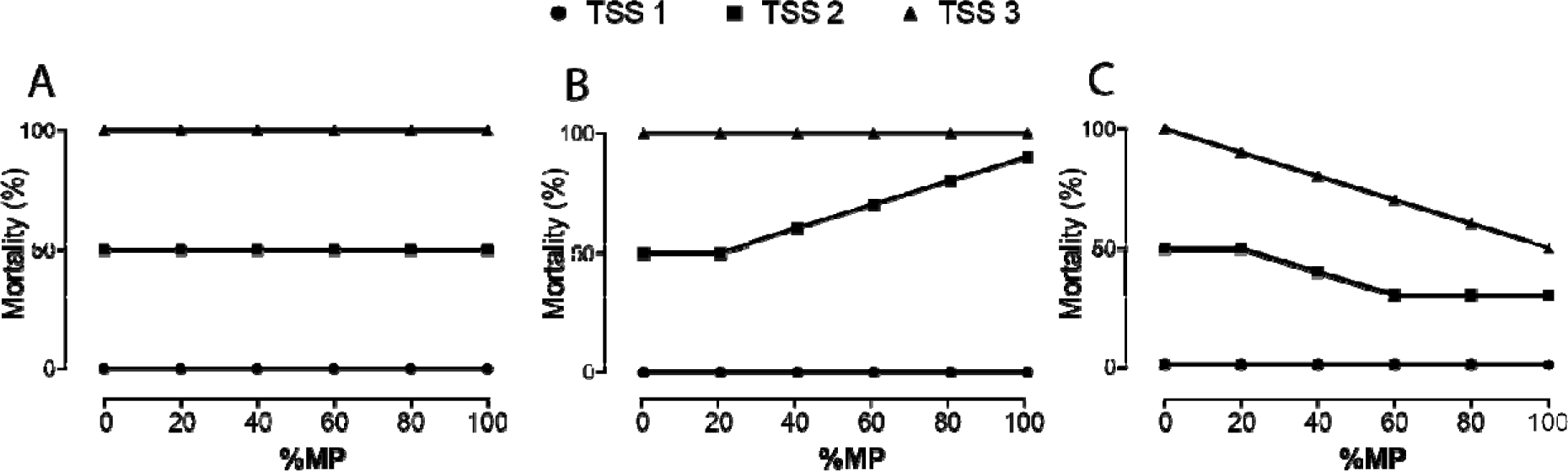
Possible outcomes of the MP Ratio Test, shown as a response (in this case Mortality%) to the relative MP contribution (%MP) to the TSS in the test system, including three test concentrations of TSS. The TSS concentrations denoted as 1, 2 and 3 represent increasing levels of TSS in the system. Scenario A shows no MP effect because mortality is only responding to the increasing TSS level. Scenario B shows an additive effect of MP because Mortality% is positively affected %MP above a critical threshold. Scenario C shows an ameliorating effect of %MP on TSS toxicity. Based on existing reports for TSS effects on *Daphnia*, we expect no effect at low TSS concentrations.

### Test organism

The freshwater cladoceran *Daphnia magna* was used as the test species. These microcrustaceans are the most common model organisms in aquatic ecotoxicology and have been used extensively in studies assessing both TSS^13,22,32–34^ and MP^13,33,35^ effects. All experimental animals originated from the same clone (Clone 5; The Federal Environment Agency, Berlin, Germany) cultured in M7 media at a density ∼10 ind. L^-1^ and fed *ad libitum* with a mixture of *Pseudokirchneriella subcapitata* and *Scenedesmus spicatus*.

### The reference and test particles

Kaolin (Sigma-Aldrich) was used as the reference particle; it occurs globally in suspended particulates and has previously been used in tests with daphnids, both as a reference particle when assessing MP effects^13^ and as a test particle when assessing effects of TSS^32^. As the test MP, we used polyethylene terephthalate (PET Goodfellow), to represent a plastic that is commonly found in the environment. The PET was obtained as 3-5 mm-sized pellets from the manufacturer and milled to a powder by Messer group GmbH, Germany. The powder was first mixed with milliQ water containing 0.01 % v/v of a non-ionic surfactant (Tween 80, Sigma-Aldrich), and sequentially wet-sieved to produce a size fraction similar to that of kaolin (for particle size distributions and details on preparation of the test particles see Fig. S1-2, Supporting Information).

### Test suspensions

Particle stocks were prepared by suspending weighed kaolin and MP with M7 media (reconstituted lake water)^36^; the volumes were adjusted to produce equal mass-based concentrations (0.1, 1, 10, 100, 1000, 10000 mg L^-1^). The test suspensions were prepared in batches with 0 %, 20 %, 40 %, 60 %, 80 % and 100 % of MP contribution to TSS. These test suspensions were then transferred to 50-mL polypropylene centrifuge tubes and used in the exposure system.

#### Experimental setup and procedures

To demonstrate the application of the MP Ratio Test, we conducted the *Daphnia* sp. acute immobilisation test (OECD 202), with some modifications. The standard *Daphnia* immobilisation test assesses 48-h mortality (immobilisation) in individuals exposed to a range of test concentrations. Based on our pilot experiments with kaolin and the reported data for MP effects on *D. magna* mortality^37^, we prolonged the test duration to 96 h. Ten daphnids (<24 h old) were placed in each test tube with the exposure media; 24 treatments (%MP × TSS concentration) were used with four replicates per treatment and two particle-free controls per run. Three TSS concentrations were tested for the 20-80 %MP mixtures and six concentrations with the single particle exposure. When sealing the test tubes, care was taken to avoid trapping air bubbles inside the tube. The tubes were mounted on a plankton wheel in a thermo-constant room at 21°C with a light: dark cycle of 16:8 h and the test was terminated after 96 h by counting live and dead animals. The daphnids were considered dead if they did not move for 30 s after being agitated.

#### Statistical analyses

For each mixture, the LC_50_ values and the corresponding 95 % confidence intervals were calculated using a dose-response curve fitted with three-parameter logistic regression (GraphPad Prism, v. 7.0; GraphPad Software, La Jolla California USA). The median mortality in the particle-free control was used as the bottom constraint and ≤100 % mortality as the top constraint. The LC_50_ values were compared across the test mixtures to evaluate the effect of %MP in the mixture; the non-overlapping confidence intervals were used as evidence of the significant difference between the treatments (%MP). Further, we estimated the corresponding LC_10_ values and used them as a surrogate for NOEC.^38^ One-phase exponential decay function was fitted to describe the relationship between the LC_10_ values and %MP in the mixture. As NOEC, we used the interpolated LC_10_ value corresponding to the lower bound of the 95 %-confidence interval for LC_10_ in pure kaolin. The critical threshold for %MP in the mixture representing no effect level of %MP in the test system was termed NOE%MP (No Effect Percentage of Microplastics).

## RESULTS

Survival in the particle-free controls was high (average 95.4 %), and mortality in relation to TSS followed the expected concentration-dependent response (Fig. S4, Supporting Information). The LC_50_ values in the treatments with 20-100 % MP were significantly lower than in the 100 % kaolin treatment, 13-43 mg L^-1^ and 482 mg L^-1^, respectively (Fig. 2), with the non-overlapping confidence intervals between the treatments with pure kaolin and the mixtures. The highest mortality was observed in the 80 % MP treatment; however, due to the broad confidence intervals for the LC_50_ values, particularly in the mixtures, the differences across all treatments with MP were not statistically significant.

**Figure 2.**
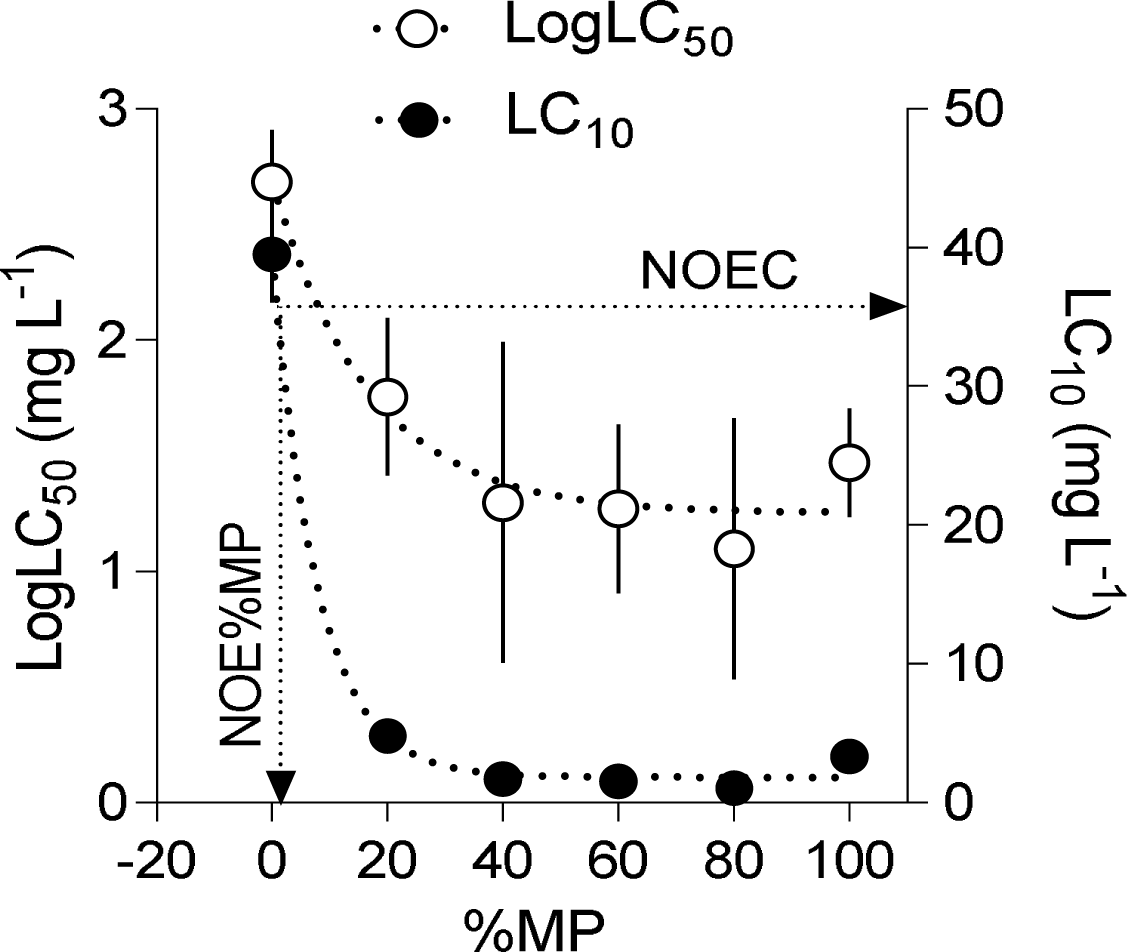
Relationships between the estimated LogLC_50_ values (mean and 95 % confidence interval; left Y-axis) and the corresponding LC_10_ values (right Y-axis) for *D. magna* and the mass contribution of MP (%MP, 0 % to 100 %) in the test mixtures of MP and kaolin. One-phase exponential decay was used to find the LC_10_ for the MP-kaolin mixture corresponding to the lower bound of the confidence interval for the kaolin treatment (NOEC) and the NOE%MP representing no effect level of %MP in the test system.

The one-phase exponential decay function provided an adequate fit for the relationship between the LC values and the %MP in the mixture (LC_50_: R^2^ = 0.95 and LC_10_: R^2^ = 0.99). The curves suggested that at %MP exceeding 40 %, the LC values levelled off at 18 and 2 mg L^-1^ for LC_50_ and LC_10,_ respectively. The fit for the LC_10_ values was used to derive the %MP threshold (NOE%MP) above which significantly higher mortality was observed. This value was determined as 1 % MP in the mixture with 36 mg L^-1^ TSS.

## DISCUSSION

### Test performance

Using the MP Ratio Test, we delineated effects of MP from those of inorganic particles of similar size and shape in the mixtures mimicking natural suspended solids in the aquatic environment. Moreover, the test design allowed estimating critical levels for the suspended MP that can be considered as hazardous. The hazardous level of MP contribution to TSS is an important outcome of our test approach because the effects of particulate contaminants, such as MP, depend not only on the absolute concentration but also on their relative contribution to the suspended matter or sediment; the latter was also recently shown by Redondo-Hasselerharm and co-workers.^39^

Ecotoxicological data describing effect thresholds in ecologically meaningful settings are needed in microplastic research to support the hazard assessment of solid polymer particles. The MP Ratio Test can be used as a tool for screening a variety of polymer materials and particle types as well as for selecting suitable test organisms and endpoints. It is conceptually similar to both the already standard bioassay approach for testing toxic effluents and sediments by serial dilutions^40^ and evaluating algal toxicity in mixtures with varying proportion of the test species.^41^ Furthermore, the need for well-characterised reference materials when evaluating, for example, particle size effects, is also recognised in nanomaterial toxicity assessment.^42,43^ However, to our knowledge, our study is the first to assess the effects of suspended MP in mixtures with natural particles using a dose-response approach.

### PET toxicity estimates

We found PET MP to be more hazardous than kaolin for *D. magna*. The addition of PET powder to the kaolin suspension increased *Daphnia* mortality, with LC_50_ values dropping more than 8-fold in mixtures with >20 % of MP. Moreover, TSS containing 1 % PET was predicted to have significantly lower LC_10_ than pure kaolin suspension. The corresponding concentration for 1 % PET-kaolin mixture was 36 mg L^-1^, which would represent NOEC of TSS containing plastics (Fig. 2).

The reported effect concentrations of MP are highly variable and span orders of magnitude even within the same level of biological organisation.^27^ Unfortunately, only a few reports provide dose-response data for MP-exposed microcrustaceans,^33,37,44^ and reference particles are rarely employed.^13,33^ Although truly comparable published data on PET toxicity for daphnids are not available; some reports are still relevant. For example, a 6-day static exposure of the copepod *Parvocalanus crassirostris* to 14 mg L^-1^ of PET (<11 µm; assuming that particles are 10-µm and spherical, with a density of 1.38 g cm³) was found to decrease population size.^45^ This concentration is four times higher compared our LC_10_ for 100 % MP. The difference, at least partially, can be explained by the fact that our animals were starved during the exposure. Moreover, the suspended amount of PET in the study with *P. crassirostris*^45^ is uncertain, because of the static exposure and the lack of information on the MP mass in the system. For other planktonic filter-feeders and various MP, the reported LC_50_ values are similar to what we have found for PET using the MP Ratio Test. For example, a 96-h LC_50_ of 57.43 mg L^-1^ was reported for *D. magna* neonates exposed to 1-µm polyethylene (PE) particles, although some mortality was observed already at 12.5 mg L^-1^.^37^ Also with *D. magna*, exposure to PE fragments (10-75 µm) produced a 48-h LC_50_ of 65 mg L^-1^.^44^ These findings are comparable with the 96-h LC_50_ observed in the 100 % MP treatment (30 mg L^-1^) as well as in the mixtures (13-43 mg L^-1^), although our values are consistently lower, which could be related to possible variations in MP aggregation and settling during the exposure as well as in specific properties of polymers and particle size.^46^ The differences in the experimental setup highlight the importance of both particle characterisation in MP research^28^ and keeping the test particle in suspension when using pelagic feeders. Still, based on either modelled data or some of the highest levels reported in the ocean,^7,47^ the levels of MP accessible for zooplankton in nature are much – approximately two to four orders of magnitude – lower than the experimentally determined MP levels for the observed effects.

### Important issues in experimental design

The straightforward logistics of our approach makes it possible to examine a large number of treatments (e.g., TSS concentrations, plastic material, particle size and shape) with reasonable effort. When testing the PET effects on the daphnid survivorship, one person was able to handle 50 experimental units per day routinely. Further developments should include a higher number of treatments with lower %MP to provide more ecologically meaningful test suspensions and improve the threshold estimates for the hazardous MP levels. Furthermore, when more information of the environmentally realistic exposure becomes available, future experimental designs can focus on narrowing concentration ranges, as well as including endpoints that are more sensitive. When testing PET, for example, a higher resolution of the %MP at the lower range (<20 %) would have provided a more precise estimate of NOEC and the corresponding %MP in the mixture. Also, various reference particles can be used depending on the research context, both natural and anthropogenic.

The selection of the reference particles is not a trivial task. One possible criterion is size because size spectra for MP and many naturally occurring particles overlap (clay: <2 μm, silt: 2–50 μm, and sand: 50 μm –2 mm).^48^ Another recently advocated option is to benchmark the test particles to the reference MP using reference plastics supplied by commercial vendors,^28^ which have a narrow size range and reliable certificates of analysis. Such reference materials would add credibility to any adverse effects exerted by unknown materials. Here, we used kaolin, because it has been relatively well studied in hazard assessment of suspended solids.^49^ Several studies also suggest its low toxicity for daphnids,^13,50^ which was supported by our 96-h LC_50_ of 482 mg L^-1^. However, in chronic tests with *D. magna*, Robinson and colleagues observed a 7-d LC_50_ of 74.5 mg L^-1,32^ which can be related to both delayed effects and possible difference in kaolin composition and aggregation during the test. Kaolin powder is a commercially available standard product, but depending on the vendor, it might contain impurities and vary in particle size distribution.

When testing effects of suspensions on planktonic organisms, it is essential to prevent that particles sediment or float, because it will affect the encounter rate and intake by the animals. Moreover, particles with different specific gravity will settle with different rates; hence performing tests under static conditions would not provide stable exposure levels. The exposure of planktonic organisms should preferably be conducted using a plankton wheel that keeps particles in suspension, thus, ensuring stable exposure conditions.^51,52^ In plankton ecology, the use of a plankton wheel is a standard procedure when conducting grazing experiments because it minimises the sedimentation of algae. Although less common in ecotoxicological testing, the plankton wheel has been used to assess the effects of suspended clay and other particulate materials on planktonic filtrators.^53^ Even though this method requires some additional effort compared to static exposures commonly employed in OECD-tests for soluble chemicals, it is a necessity for standardising the exposure conditions.

Another area of concern with respect to the standardisation of the testing procedure is particle behaviour, such as aggregation and settling, during the exposure. Since suspended particles may have complex behaviour, their dispersion stability during exposure should be controlled. The extent of aggregation, i.e., aggregate size, composition and cohesion, is dependent on the complex interplay between various components, such as particle material, size, concentration, media ion composition, and organic material present. Aggregation may have affected the amount and size spectra of PET and kaolin and could potentially explain the non-linear response of LC to %MP. The relationship between the aggregate formation and the measured response is particularly relevant at high TSS concentrations. On the other hand, aggregation, at least to some extent, is preventable by, for example, dispersants and sonication, which may be sufficient in short-term experiments. These methods are employed in nanomaterial effect assessments^42^ and, in some cases, microplastic studies.^54^

The exposure duration is yet another issue that needs to be considered in acute tests. Daphnid energy expenditure may increase in the presence of non-food particles since the induced filtering activity is similar as for food particles,^55^ while the cost of cleaning appendages and egestion through postabdominal rejections increase.^15^ Sensitivity to the refractory material would increase with starvation, and the higher energy expenditure may decrease survival. In *D. magna* neonates, the critical exposure time is 96 h, which marks the depletion of their fat reserves.^56^ Earlier studies using the *D. magna* acute immobilisation test for particle suspensions have also shown that it is suitable to extend the exposure period to 96 h from the standard 48 h to increase the sensitivity of the test.^37^ However, if a test organism other than *Daphnia* neonates is used, the duration of the exposure must be adjusted depending on its capacity to withstand starvation.

### Implications for regulatory measures and concluding remarks

Many suspension- and filter-feeders frequently face turbid environments with high concentrations of refractory materials generated by natural processes, such as terrestrial runoff, currents, and weather-induced bottom sediment resuspension,^57^ but also by anthropogenic activities, such as dredging and capping of contaminated sediments using various materials. In aquatic ecology, conditions with elevated TSS are acknowledged as stressful^58^ and regulated by water quality standards.^24,59^ The quality standards vary across regions and types of aquatic systems, e.g., lotic and lentic, and systems with different natural levels of suspended solids. For example, the Alaskan state standard for clear-water lakes is a maximum increase of TSS, above background levels, equivalent to 25 mg L^-1^, whereas an increase of 100 mg L^-1^ is acceptable for streams.^25^ Similarly, hazard assessment of MP in different systems would eventually require that effect-thresholds are established for the critical MP concentration, such as NOE%MP, relative to the background TSS levels and their combined interactions with biota.

By using reference particles – natural minerals or standardised plastics – with predictable effects on the test organisms, one may identify and account for the general responses anticipated from exposure to suspended solids. The relative importance of MP addition to the suspension could thereby be assessed. Interestingly, the hazard level we found for MP contribution to TSS using a planktonic organism is virtually identical to the EC_10_ of 1 % MP per sediment dry weight, reported for a benthic macroinvertebrate.^39^ This could further support the possibility to benchmark MP effects against the lower 95 % confidence bound of the reference material and using the corresponding contribution of MP in the mixture to estimate the hazard level of MP (Fig. 2).

The hazard assessment and the regulatory framework for MP contaminants in aquatic systems require integration with an assessment of particulate matter pollution at large because the approaches required to establish toxicity are similar. Moreover, raising levels of black carbon in the atmosphere implies increased inputs of these particles in the aquatic systems, where their environmental effects are also a matter of concern.^60^ Addressing all types of particulate pollution and focusing on physicochemical properties of these particles would provide a translational value when developing testing and regulatory practices.

## ACKNOWLEDGEMENT

This research was funded by projects WEATHER-MIC, irPLAST and MICROPOLL, which are supported through the Joint Programming Initiative Healthy and Productive Seas and Oceans (JPI-Oceans), Swedish Research Council for Environment, Agricultural Sciences and Spatial Planning (FORMAS), the joint Baltic Sea research and development programme (BONUS) and the Swedish Innovation Agency VINNOVA.

**Supporting Information.** Additional text and four figures describing particle preparation and size distributions analyses, confirmed particle ingestion and separate dose-response curves for all the tested mixtures of and kaolin and PET.

